# Ocean currents promote rare species diversity in protists

**DOI:** 10.1101/2020.01.10.901165

**Authors:** Paula Villa Martín, Ales Bucek, Tom Bourguignon, Simone Pigolotti

## Abstract

Oceans host communities of plankton composed of relatively few abundant species and many rare species. The number of rare protists species in these communities, as estimated in metagenomic studies, decays as a steep power law of their abundance. The ecological factors at the origin of this pattern remain elusive. We propose that oceanic currents affect biodiversity patterns of rare species. To test this hypothesis, we introduce a spatially-explicit coalescence model able to reconstruct the species diversity in a sample of water. Our model predicts, in the presence of oceanic currents, a steeper power law decay of the species abundance distribution and a steeper increase of the number of observed species with sample size. A comparison of two metagenomic studies of planktonic protist communities in oceans and in lakes quantitatively confirms our prediction. Our results support that oceanic currents positively impact the diversity of rare aquatic microbes.

## Introduction

Oceanic plankton can be transported across very large distances by currents. Many planktonic species are cosmopolitan, i.e. they are found virtually everywhere across the global ocean (*1, 2*). These observations suggest that, at first sight, the distribution of planktonic species is not limited by dispersal, and therefore that niche preference is the predominant factor determining species abundance (*3*). However, niche theory predicts species-poor planktonic communities for plankton thriving on a limited set of resources. Observations contradict this prediction and show that planktonic communities are very diverse (*4–7*). This violation of the basic principles of niche theory (*8, 9*) has puzzled ecologists for decades (*10*) and has fostered numerous studies attempting to explain the diversity of plankton (*11*). One proposal is that variable environments offer more possibilities for specialization of ecological traits (*5, 12–18*). Another proposal is that oceanic currents can create barriers reducing competition among species, and therefore promoting species coexistence (*19, 20*). Quantitative analyses also suggest that oceanic currents play a significant role in organizing large-scale diversity patterns (*21, 22*), and that dispersal limitation contributes, alongside with niche specialization, to microbial biodiversity of oceans (*23–27*).

DNA metabarcoding has allowed rapid and extensive measurements of the diversity of aquatic microbial communities, providing new means to study the ecological forces shaping planktonic communities. Metabarcoding studies have revealed that, beside commonly observed species, planktonic communities are characterized by a vast range of rare species. This so-called “rare biosphere” (*28, 29*) makes up the majority of planktonic species (*25, 30*). The diversity of planktonic species can be quantified by the Species Abundance Distribution (SAD), defined as the frequency *P*(*n*) of species with abundance *n* in a sample. SADs of rare marine protists are qualitatively different from those of abundant species (*31, 32*) and appears to follow a power law distribution

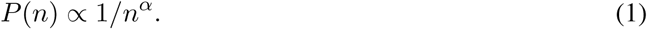

The exponent *α* varies significantly among samples, appears weakly correlated with environmental factors, and is significantly larger than 1 on average (*33*). Diversity patterns in other microbial communities, such as that of the human gut (*34*), are well described by a form of SAD following the Fisher log series, *P*(*n*) ∝ *e*^−*cn*^*/n* (*35–37*), as predicted by Hubbell’s neutral model (*36, 38, 39*). For large communities, the parameter *c* is very small, so that the distribution is close to a power law with *α* = 1. Hubbell’s neutral model is therefore unable to explain the decay of SADs in oceanic protist communities. Steep SADs, in agreement with the data, can be obtained with a modified neutral model that takes into account density-dependence of growth and death rates (*33, 40–42*). However, the ecological forces determining this density-dependence in the oceans are unknown.

In this paper, we propose that the steep decay of SADs observed in the oceans is caused by the particular way oceanic currents limit dispersal. Although several studies have shown that currents can affect effective population size (*43*) and provoke counterintuitive effects on fixation times (*44–46*), these studies did not scrutinize the effect of currents on multi-species communities. To test our hypothesis, we introduce a model that takes into account the role of oceanic currents in determining the genealogy of microbes in a sample. Our model predicts that, in the presence of oceanic currents, SADs are characterized by larger values of the exponent *α*. We also predicts that currents cause a sharper increase of species diversity as a function of sample size. To test these predictions, we analyze 18S rRNA sequencing data generated from oceanic (*33*) and lake protist communities (*47*). The observations quantitatively match our predictions, supporting the idea that oceanic currents are responsible of the differences in biodiversity patterns between oceans and lakes.

## Results

### Coalescence model predicts the effect of oceanic currents on SADs

We introduce a computational model to assess the effect of oceanic currents on the protist species distribution of a water sample. In this model, we assign a tracer with spatial coordinates to each individual in the sample (see Fig. 1). Tracers are initially placed in a local area, representing the portion of water where the sample was collected. The coordinates of each tracer move backward in time, following the spatial trajectory of the ancestors of each individual (see Fig. 1). If two tracers are at a sufficiently close distance, they coalesce into a single tracer with a given probability. This new tracer represents the most recent common ancestor of the two individuals. Finally, tracers are assigned at a fixed rate *µ* to one species. These events represent immigration due to other causes than ocean currents; assigned tracers are eliminated from the system. At the end of the run, individuals in the original sample are considered conspecific if they have coalesced to a common ancestor before being eliminated (see Fig. 1 and Methods). This coalescence model can be interpreted as the backward version of an individual-based community model which includes the effect of fluid flows (*44, 48*) (see Supplementary Fig. 1). The coalescent formulation has the advantage of describing the dynamics of one sample embedded in a larger ecosystem (*49, 50*).

**Figure 1:**
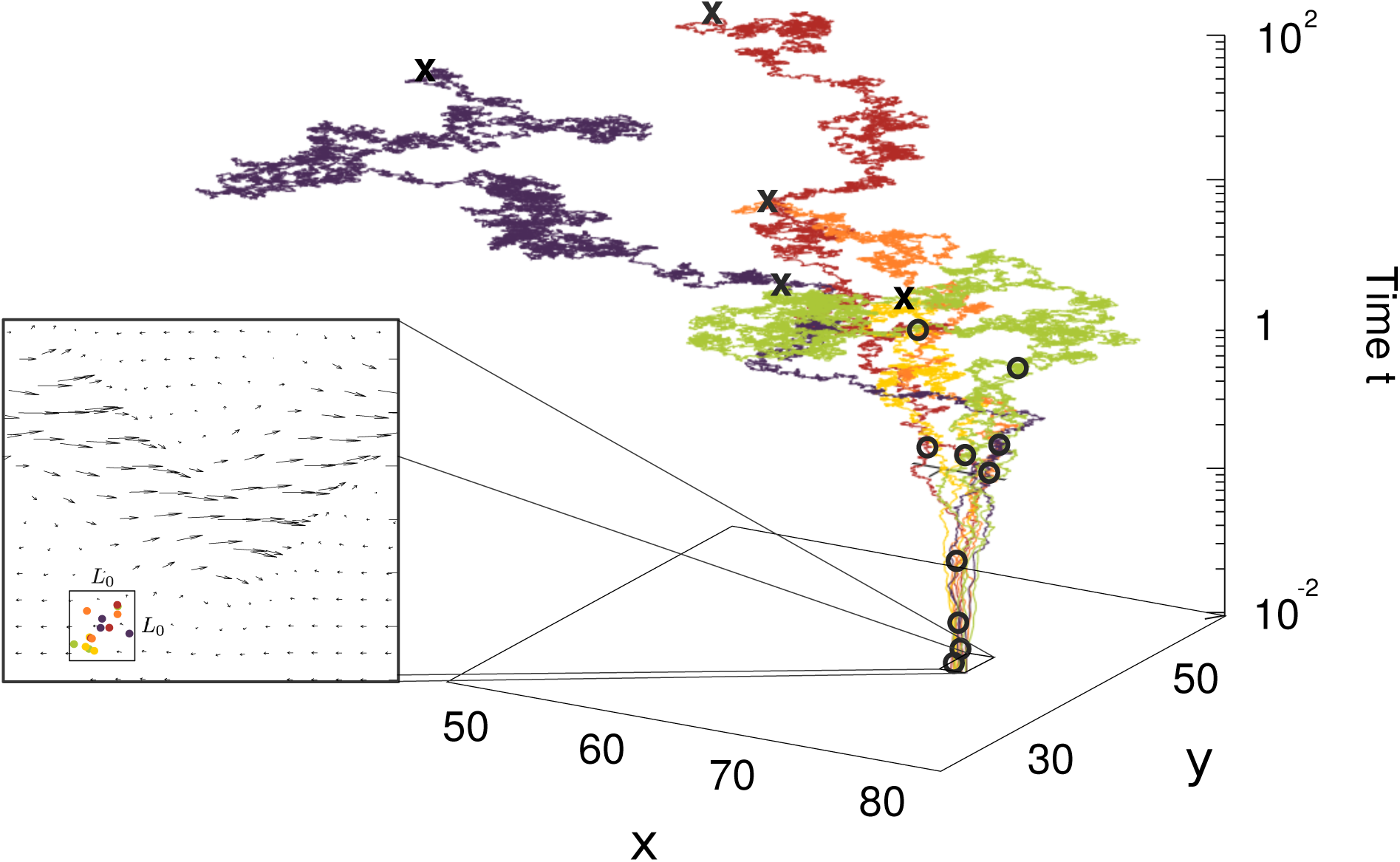
Genealogy in oceanic currents. (left) The coalescence model predicts the protist species composition of a sample of oceanic water of size *L*_0_ × *L*_0_. Different colors represent different species. Arrows represent the velocity field caused by ocean currents. (right) Trajectories of the coalescence model with ocean currents. Individuals are represented by tracers, that are transported backward in time and can coalesce with other tracers if they reach a close distance. Coalescence events are marked by open circles; trajectories of individuals that have coalesced are shown in the same color. Tracers are removed from the population at an immigration rate *µ* (marked by crosses). See also Supplementary video.

We simulate the coalescence model with and without oceanic currents. In the latter case, movements of tracers are modeled as a simple diffusion process, taking into account individual movements and small-scale turbulence. In the former case, we superimpose to this diffusion process the effect of large scale oceanic currents. We model oceanic currents with a kinematic model of a meandering jet, which is a ubiquitous large-scale structure characterizing oceanic flows (*51–54*). Population sizes and parameters characterizing the flow are sampled in a physically realistic range (*53, 54*) (see Methods). All other parameters characterizing population dynamics are chosen identically in the two cases (see Methods).

SADs predicted by the model display a considerable variability depending on parameters and demographic stochasticity, both in the presence and absence of currents (see Fig. 2a and 2b). To characterize individual SAD curves, we fit them with a power law function *P*(*n*) ∝ 1*/n*^*α*^ using maximum likelihood in an optimal range of abundances (see Methods). For comparison, we also fit an exponential *P*(*n*) ∝ *e*^−*cn*^, and a Fisher log series *P*(*n*) ∝ *e*^−*cn*^*/n* in the same range. The power law provides a better fit than the exponential in most cases (74% and 77% of samples with and without currents, respectively) and than the Fisher log series (75% and 62% of samples with and without currents, respectively).

**Figure 2:**
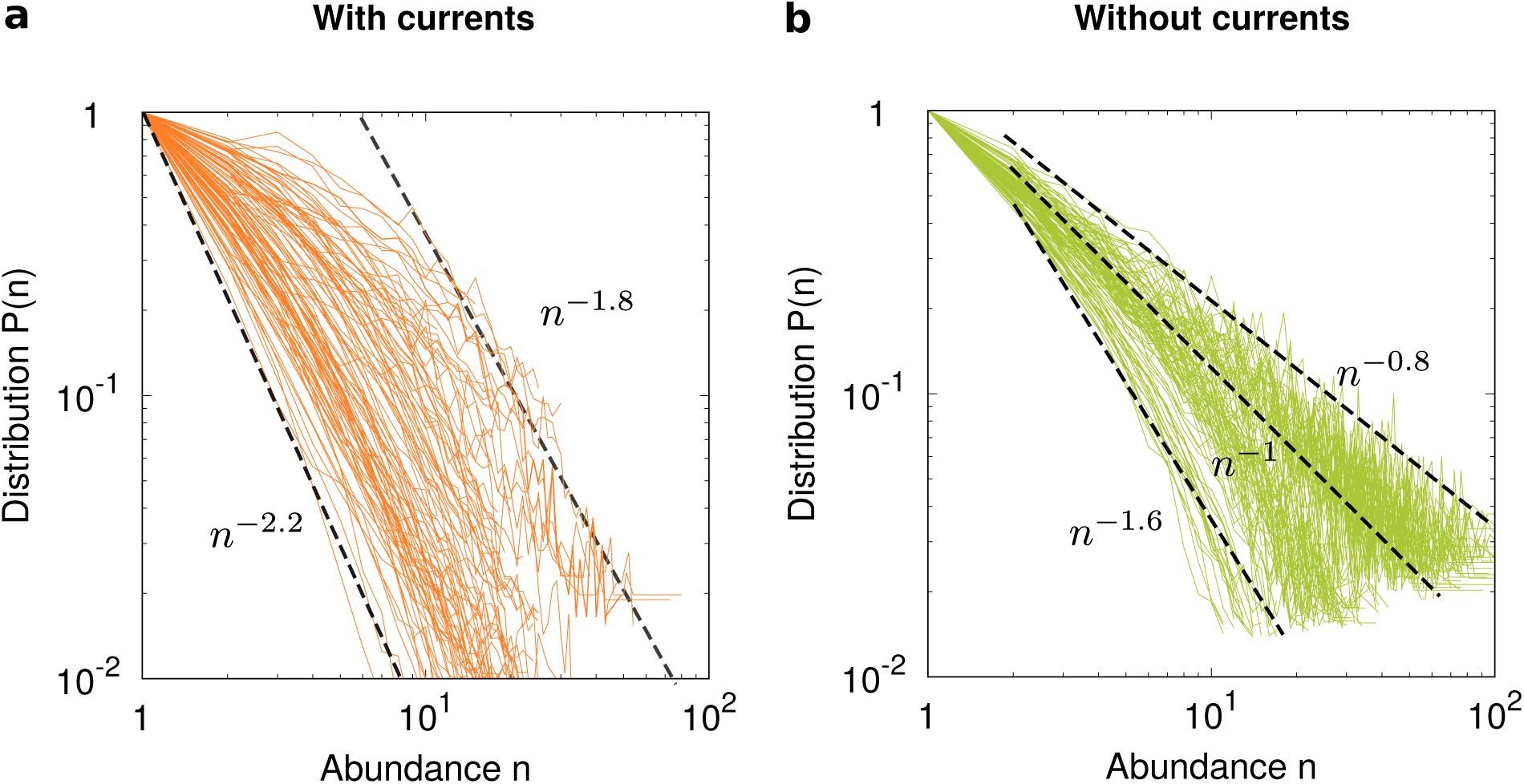
Coalescence model predicts effect of currents on species abundance distribution. We show SADs A) in presence (orange lines) and B) absence (green lines) of oceanic currents for the coalescence model. Here and in the following, SAD curves are rescaled so that *P*(1) = 1 to ease visualization. Model details and parameters are presented in Methods. Dashed lines are power laws to guide the eye (see also Fig. 3).

Introducing oceanic currents in the model increases, on average, the steepness of SADs (see Fig. 2a and 2b). We investigate the physical mechanism causing this effect. One effect of transport by currents is to enhance the effective diffusivity (*55*). We test whether this effect is responsible for the change of pattern in the SAD by running our model with the effective diffusivity of the kinematic model, but without currents (see Supplementary Fig. 2). We find that the SADs are weakly affected by the change in diffusivity. This means that the change in SADs must be due to the structures created by the oceanic currents, which can not be simplified into a diffusion process. We further run our model with a parameter choice yielding currents constant in time (see Supplementary Fig. 3). Neither in this case we observe the steep SADs as in the presence of time-dependent currents. Taken together, these results suggest that the time-varying, chaotic nature of oceanic transport is responsible for the steepening of SAD curves.

### Protist SADs are steeper in oceans than in freshwater

To test our predictions, we analyze DNA metabarcoding datasets from two studies of aquatic protists. The first dataset includes oceanic protist DNA sequences of 157 water samples from the TARA ocean expedition (*33*). The second dataset includes protist DNA sequences of 184 freshwater samples taken from lakes (*47*). In both cases, we use 97% sequence identity threshold to cluster protist sequences into operational taxonomic units (OTUs). We calculate SAD for each sample of both datasets using OTUs as proxies for species (see Methods).

From now on we discard “abundant species”, defined as those in abundance classes *P*(*n*) including less than 4 species. The remaining “rare species” are the subject of our study and constitute 93% of all species in ocean samples and 78% of all species in lake samples.

As for the model, SAD curves display considerable sample-to-sample variability, both in ocean and in freshwater samples (see Fig. 3). The variability is possibly caused by differences in size, protist community composition, and ecological conditions among samples. Observed SAD curves are better fitted by a power law than exponential or Fisher log series in most cases. The exponential distribution provides a better fit than the power law in 13% of lake samples and 13% of oceanic samples, whereas the Fisher log series provides better fits than the power law in 39% of lake samples and 18% of oceanic samples. We obtain similar results with different OTU definition (95% instead of 97% similarity) and different thresholds separating abundant from rare species (see Supplementary Table 1 and 2).

**Figure 3:**
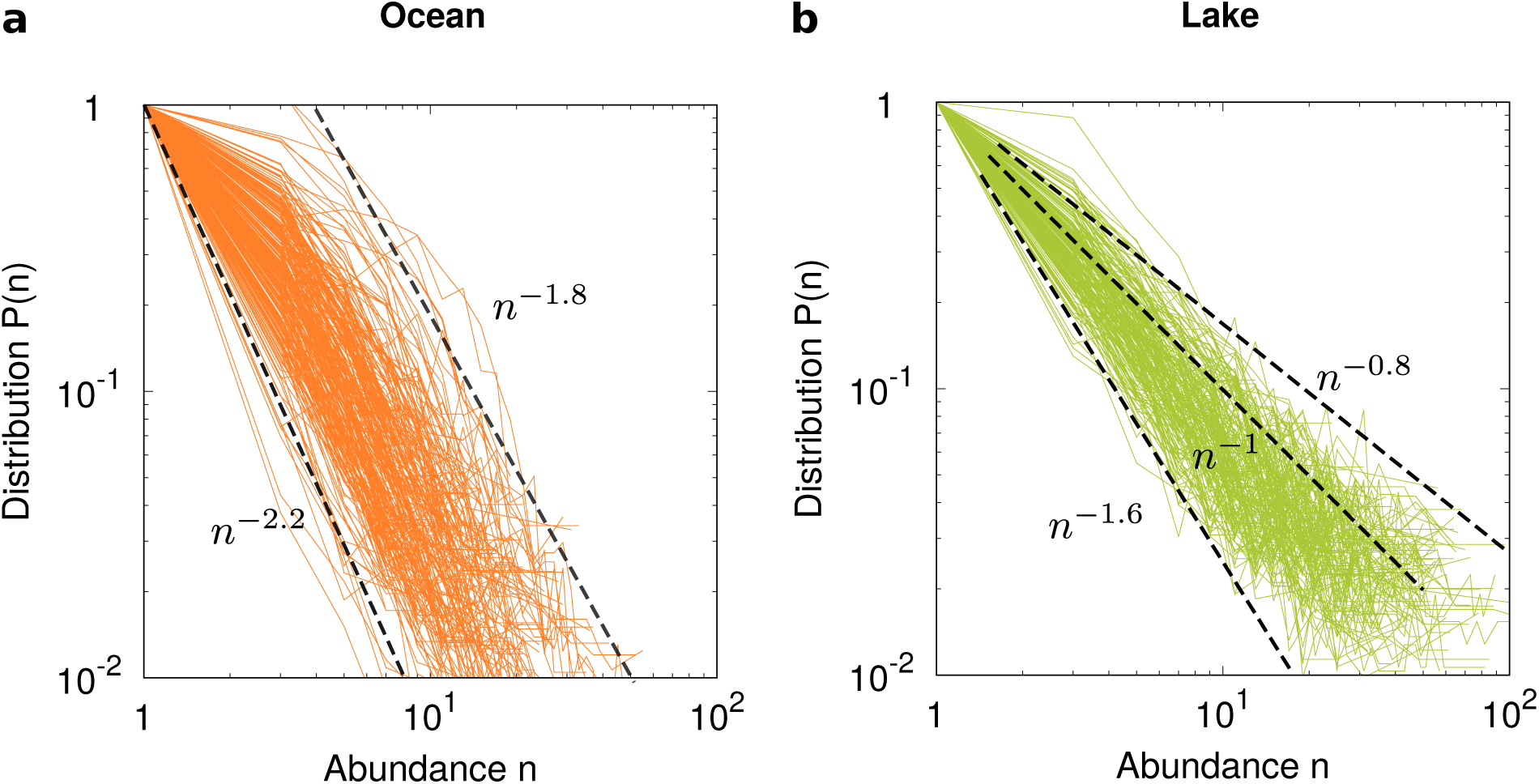
Rare species abundance distributions present a steeper decay with abundance in oceans than in lakes. Continuous lines represent SADs of protist communities from (a) 157 oceanic (*33*) and (b) 184 freshwater (*47*) samples. Total numbers of individuals in each sample are in the ranges of (a) (10^3^, 10^5^) and (b) (10^4^, 10^6^). In both panels, power laws (dashed lines) are shown to guide the eye.

Strikingly, the power law decay of SADs is on average steeper in oceans than in lakes (see Fig. 3), as predicted by our coalescence model.

### Distribution of the SAD exponent is quantitatively predicted by the coalescence model

We quantify the agreement between model and data by analyzing the distribution of the power law exponent *α* in equation (1). In the presence of currents, the model predicts a value of the exponent significantly larger than one (average *α* = 1.70, standard deviation *σ* = 0.68). In the absence of oceanic flows, the model predicts an average *α* = 1.26, (*σ* = 0.46), a value compatible with the neutral prediction *α* = 1 in well-mixed systems (*35–37*) and spatially-explicit neutral models (*50, 56*).

Observations in both oceans and lakes are in excellent agreement with the distributions of exponents predicted by the model (see Fig. 4a). Our analysis confirms that the average exponent *α* is significantly larger than 1 in the oceans (average *α* = 1.79, *σ* = 0.52, see Fig. 4a and (*33*)). In the lakes, the average exponent is *α* = 1.15 (*σ* = 0.36, see Fig. 4a). Adopting a different definition of OTUs (95% instead of 97%) and/or different thresholds separating abundant from rare species leads to qualitatively similar result (see Supplementary Tables 1 and 2 and Supplementary Fig. 4). In particular, the average exponent *α* in the oceans is between 8% and 43% larger than in the lakes, depending on the threshold and the definition of OTUs.

**Figure 4:**
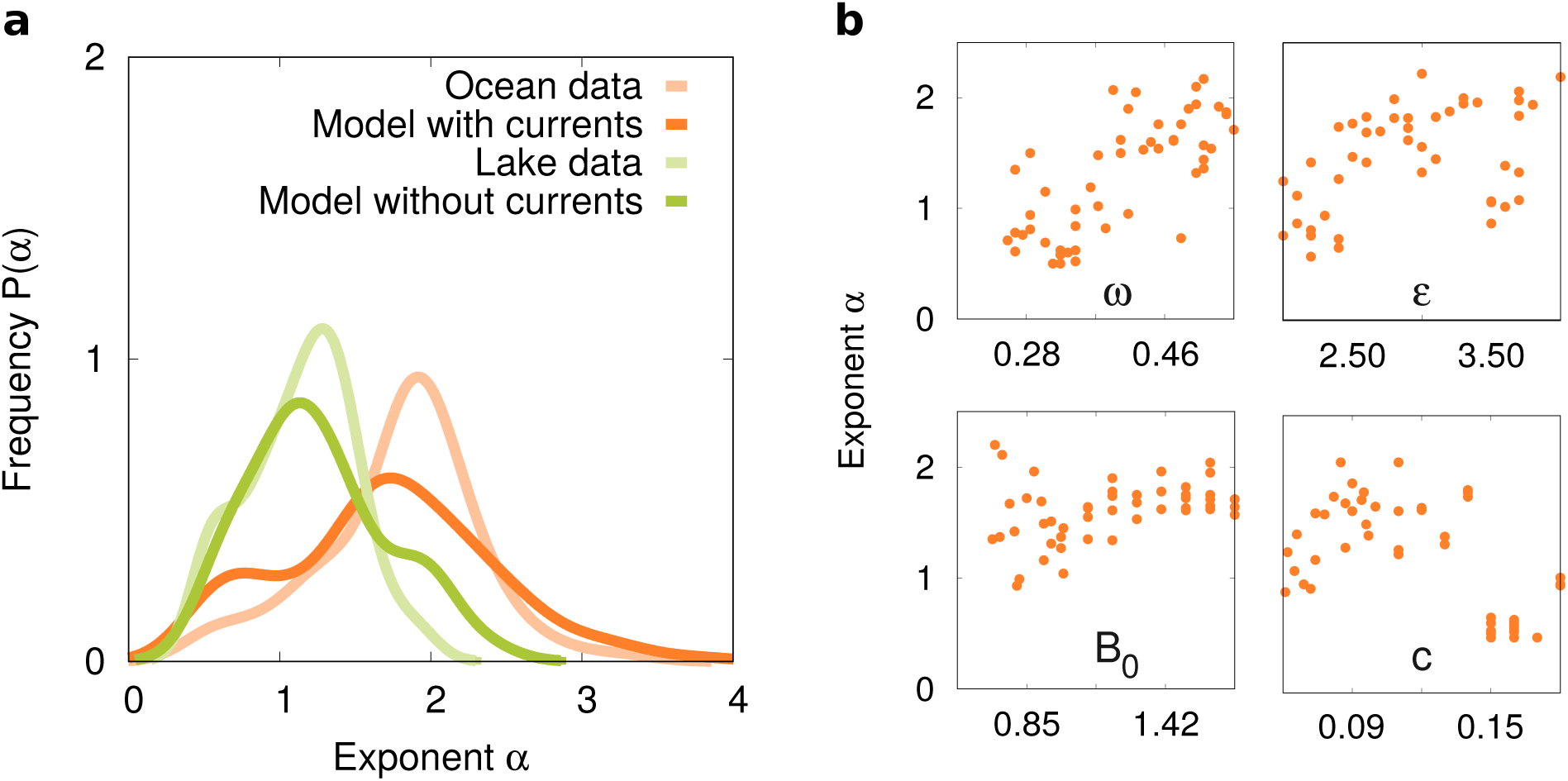
Power law exponents of species abundance distributions in the global ocean. We run our models for different population sizes and different values of flux parameters for ocean samples (see Methods). We select 157 oceanic samples and 184 freshwater samples as in Fig. 3. We fit the power law exponent *α* of the SADs to the model and to the data using maximum likelihood (*57–59*). (a) Continuous distributions of the exponent obtained by kernel density estimation (*60, 61*). (b) Dependence of the exponent on four main parameters of the oceanic flow. In each sub-panel, other parameters are kept constant (see Methods).

We find that four parameters characterizing the shape and the mixing level of the jet mostly affect *α*. However, the value of the exponent does not present a regular trend upon varying each parameter individually (see Fig. 4b and Supplementary Fig. 5).

### Ocean currents lead to a steeper increase in number of species as a function of sample size

By simulating our model at varying sample size *N* with and without currents, we predict that currents should significantly increase the number of expected species in each sample (see Fig. 5a). This effect is consistent with the increase of *α* in the presence of currents: increasing *α* suppresses very abundant species, and therefore makes the samples more diverse. This effect becomes more and more pronounced as *N* is increased. In the data, we find that samples from oceans contain more species than samples from lakes at similar sample size, consistently with our prediction (see Fig. 5a). The observed enrichment is even stronger than predicted by our model.

**Figure 5:**
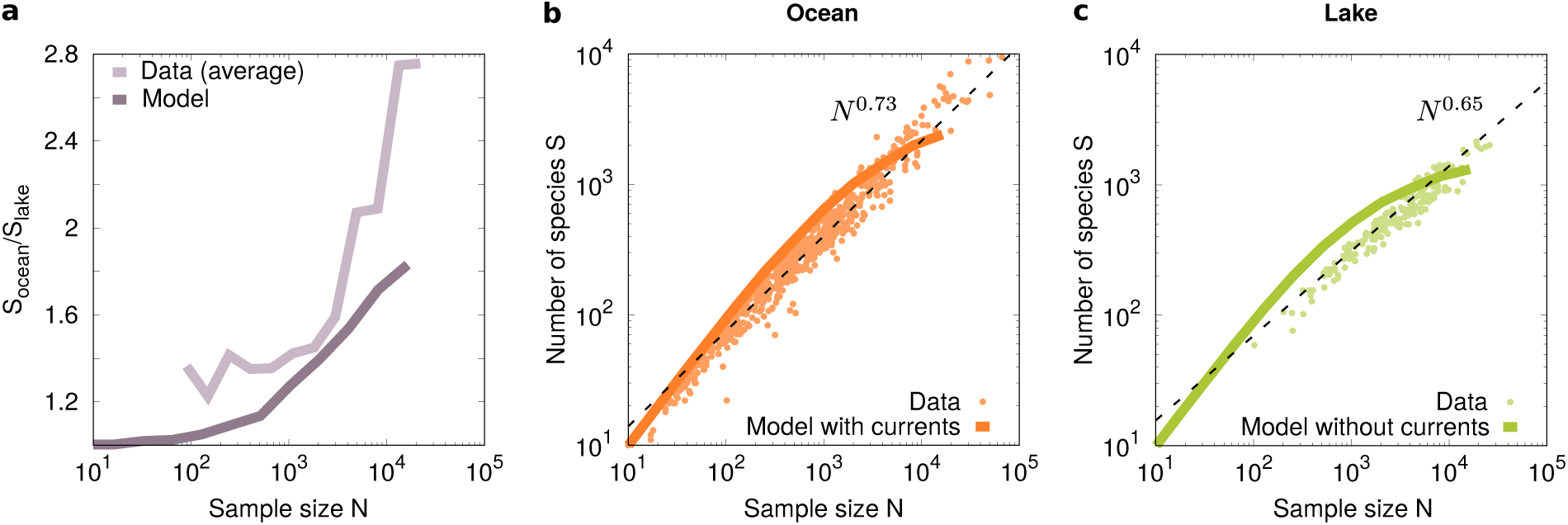
Oceanic currents increase species number *S* in a water sample. (a) Ratio *S*_ocean_*/S*_lake_ as a function of the sample size *N* for the model and the data. We simulate the model at increasing sample sizes *N* in powers of 2 and obtain continuous curves by interpolation. Other parameters are presented in Methods. Averaged data are obtained by binning for both oceans and lakes. (b,c) Number of species *S* in samples of *N* individuals in (b) oceans and (c) lakes. A power law (equation 3) fits the data better than Ewens sampling formula (equation 2) for both (b) oceanic (normalized log likelihood −19.39 versus −449.52) and (c) lake samples (normalized log likelihood −12.46 vs −111.94). Fitted exponents are *z* = 0.73 and *z* = 0.65 for oceans and lakes, respectively. The results of the coalescence model are shown with and without oceanic currents (orange and green line, respectively). Ewens sampling formula provides a better fit than the power law in both cases, (b) −420.95 vs −2131.43 and (c) −434.66 vs −1902.62.

We now study the increase of number of species with sample size in oceanic and lake water samples individually. In the case of well-mixed populations, the species composition of a given sample is described by the Ewens sampling formula (*62*), which predicts that the expected number of species in the sample is

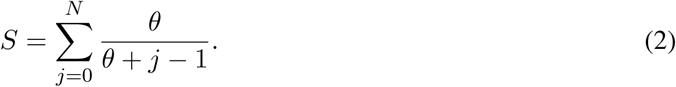

where *θ* = 2*N*_eff_*µ* is the fundamental biodiversity number (*36*) and *N*_eff_ is the effective population size. Alternatively, sample species composition can be empirically described using a power law (*63, 64*)

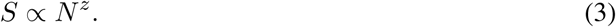

Our model predicts an increase in the number of species with the sample size largely as predicted by the Ewens sampling formula (see Fig. 5a and Fig. 5b). Both for ocean and freshwater samples, the power law model provides a better fit (see Fig. 5a and Fig. 5b) with a higher exponent for oceanic samples (*z* = 0.73) compared to lake samples (*z* = 0.65).

## Discussion

Oceanic currents are known to largely affect plankton distribution at large scale (*19–21*). Here, we show that ocean currents have profound effects on plankton distribution and diversity of rare protist species even at the level of individual metagenomic samples. Our coalescence model bridges the gap between large-scale oceanic dynamics and ecological dynamics at the individual level and provides a versatile and powerful tool to validate individual-based ecological models using DNA metabarcoding data. Although, for simplicity, we focus on neutral dynamics and protists, our approach can be extended to more general ecological settings and to other plankton communities, including animals and prokaryotes. Such generalizations, combined with high-throughput sequencing data, has the potential to shed light on the main ecological forces determining plankton dynamics.

The coalescence model predicts that oceanic currents are responsible for steeper decay of SAD curves and steeper increase in the number of observed species with sample size. Both these predictions are in quantitative agreement with observations. The steep decay of SAD distributions in the oceans has been previously explained in terms of density-dependent effects (*33*). Although our study does not preclude this possibility, the comparison with freshwater ecosystems strongly suggests that oceanic currents effectively determine this density dependence. The steeper decay of SAD curves predicted by the coalescence model depends on geophysical parameters characterizing mixing of oceanic currents. The irregular behavior of the SAD exponent as a function of these parameters (see Fig. 4b) potentially explains the difficulty of correlating observed values of *α* with other oceanographic measurements (*33*).

Oceanic flows, such as those considered here, act as barriers limiting transport. Our results support that these barriers reduces the pace of individual coalescence into species, and therefore limits the formation of abundant species. The comparison with a non-chaotic flow and with purely diffusive dynamics supports that the chaotic nature of oceanic flows is the main cause of the steep SAD exponents. A detailed physical theory of this phenomenon remains a challenge for future studies.

In summary, our study provides a mechanistic theoretical framework to analyze diversity of rare microbial species in aquatic environments at the individual level and paves a way to quantitatively understand how dispersal limitation by oceanic currents shapes the diversity of planktonic communities.

## Methods

### Coalescence model

We consider *N* microbial individuals in an aquatic environment and seek to reconstruct their ancestry. Each individual is associated to a tracer with two-dimensional spatial coordinates *x, y*. Initially, tracers are homogeneously distributed in a square *L*_0_ × *L*_0_, representing the area where the sample was collected. The tracers move in space according to the stochastic differential equations

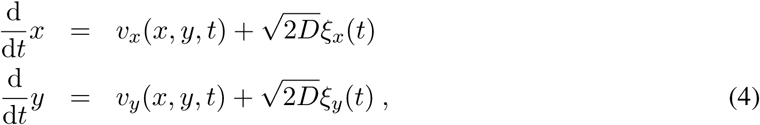

where *v*_*x*_, *v*_*y*_ is an advecting field representing the effect of oceanic currents, and the terms proportional to 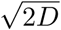 are diffusion terms modeling individual movement and small-scale turbulence. The quantities *ξ*_*x*_(*t*), *ξ*_*y*_(*t*) are independent white noise sources satisfying ⟨*ξ*_*i*_(*t*) ⟩ = 0, ⟨*ξ*_*i*_(*t*)*ξ*_*j*_(*t*′) ⟩ = *δ*_*ij*_*δ*(*t* − *t*′) where ⟨… ⟩ denotes an average and *i, j* ∈ (*x, y*). The advecting field *v*_*x*_, *v*_*y*_ is discussed in the next subsection. Since the coalescence model evolves backward in time, we integrate equation (4) from the final configuration with negative time increments.

Tracers at a short distance *δ* from each other can coalesce at a rate *r*. We implement immigration events by assigning species at a rate *µ*. At each time step d*t*:

1. All tracers move from (*x, y*) to (*x* + Δ*x, y* + Δ*y*) by numerically integrating equation (4).
2. Tracers are selected one by one and are removed with probability *µ* d*t* (immigration event). Tracers coalesce with probability *r* d*t* when an individual *j* is in a area of size *δ* × *δ* centered at the selected individual *i*.

We set *r* = 1, *µ* = 10^−4^, and the diffusion constant to *D* = 3 · 10^−9^ as further discussed below. The interaction distance *δ* is chosen to satisfy *D* = *rδ*^2^, see (*48*). We consider sample areas of linear size on the order of the mean distance traveled by an individual in one generation, *L*_0_ = 5 Km, estimating a protist life time of about one day (*65*), and protist movements of about 20 Km^2^ per day (*66*). Population size is randomly selected for each run in the range *N* ∈ (10^3^, 10^5^) unless otherwise indicated. For Fig. 4b and Supplementary video we set *N* = 8192.

### Kinematic model of the oceans

We model large-scale oceanic currents by means of a kinematic model of a meandering jet (*51–54*). The velocity field *v*_*x*_, *v*_*y*_ is defined in terms of a stream function. In a fixed reference frame, the stream function reads

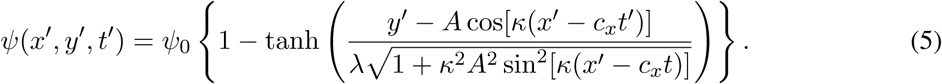

The stream function is more conveniently written in a dimensionless form

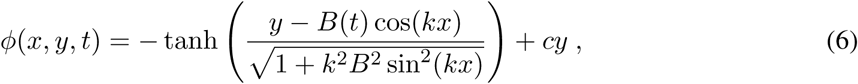

being *B*(*t*) = *B*_0_ + *ϵ* cos(*ωt* + Φ), *c* = *c*_*x*_*L/ψ*_0_ and *k* = 2*π/L* with *L* the meander wave-length. The transformation between dimensional and dimensionless units is *x* = *x*′*/λ, y* = *y*′*/λ* and *t* = *t*′*ψ*_0_*/λ*^2^ (*52*). Given the stream function, the components of the velocity field in dimensionless units are

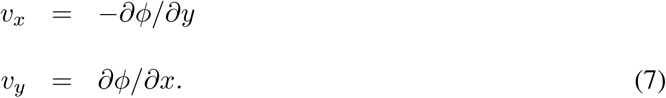

We run the simulations in a virtually infinite system. In the case without currents, this modeling choice is justified a posteriori by the fact that, based on our observations, lake SAD exponents do not present a significant dependence on lake area (see Supplementary Fig. 6). For the ocean simulations, results can be affected by the position of the local area. For this reason, the sample area *L*_0_ × *L*_0_ is placed at random coordinates *x*_0_, *y*_0_ ∈ (0, 8) for each run. For Fig. 5 we fix *x*_0_ = 7.5 and *y*_0_ = 1.

### Parameters of the kinematic model

Realistic parameters of the dimensionless stream function, equation (6), are estimated as *L* = 7.5, *c* = 0.12, *B*_0_ = 1.2, *ω* = 0.4, *ϵ* = 0.3 and, Φ = *π/*2 (*53, 54*). We consider parameter ranges based on these values *c* ∈ (0.06, 0.18), *B*_0_ ∈ (0.7, 1.7), *ω* ∈ (0.25, 0.55) and fix *L* = 7.5, Φ = *π/*2. The value of *E* has to be larger than a critical value depending on *ω* to prevent transported particles to remain trapped into long-lived eddies (*54*). To meet this condition while exploring a range of values of *ω*, we fix *ϵ* ∈ (2, 4). For Fig. 4b, Fig. 5 we set *c* = 0.12, *B*_0_ = 1.2, *ω* = 0.5, and *ϵ* = 3.

To convert from dimensionless units to dimensional units, we use the spatial scale *λ* = 40 Km (*51*) and the stream function scale *ψ*_0_ = 160 Km^2^ day^−1^. With this choice, the time unit *λ*^2^*/ψ*_0_ is equal to one day. The parameter *ψ*_0_*/λ* represents the maximum velocity in the center of the jet. With our choice of units, the velocity is equal to 40 Km day^−1^, slightly lower than the average velocity of large-scale oceanic currents (about *ψ*_0_*/λ* ≈ 200 Km day^−1^ for the surface Gulf stream and 50 Km day^−1^ for the lower thermocline (*51*)).

In physical units, the coalescence rate is equal to *r* = 1 day^−1^, i.e. about one generation time for protists (*65*). Our choice of the diffusion constant to *D* = 3 · 10^−9^ in dimensionless units corresponds to about 6 · 10^−5^*m*^2^*/s* in physical units, which is consistent with observations (*55*).

### Species abundance distribution (SAD)

We compute the distribution *P*(*n*) of the species abundances *n* for each sample. Species with low to intermediate abundance appear to follow different distribution than abundant species, as previously observed (*31–33*). For this reason, we filter out species in abundance classes below *P*(*n*) = 4. To avoid overfitting, we also discard samples with SAD composed of less than 10 points with different frequencies *P*(*n*). After this selection, we are left with 157 samples for oceans and 184 for lakes. We compute the species abundance distribution *P*(*n*) for the coalescent model with and without advection. Each sample is obtained for different flux parameters and population sizes (described above). The resulting distributions *P*(*n*) are averaged over up to 10^2^ realizations of the model and filtered in the same way as the data samples for consistency.

### Data fits

To determine the exponent *α*, we fit the function

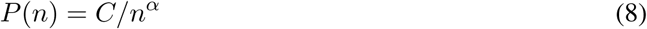

in a range of intermediate abundances (*n*_min_, *n*_max_). The exponent *α*, the proportionality constant *C*, and the values of *n*_min_ and *n*_max_ are simultaneously determined by maximizing the normalized log-likelihood ln *L* = (1*/*𝒩) ∑_*i*_[*n*_*i*_ ln *P*(*n*_*i*_) − *P*(*n*_*i*_)+ln(*n*_*i*_!)], where 𝒩 is the number of non-zero abundance classes for *n* in the range (*n*_min_, *n*_max_) and we assumed Poissonian counts (*57–59*). We discard samples for which the range (*n*_min_, *n*_max_) includes less than 5 points. We also fit an exponential *P*(*n*) = *Ce*^−*cn*^ and a Fisher log series *P*(*n*) = *Ce*^−*cn*^*/n* with the same method and in the same interval [*n*_min_, *n*_max_] determined with the power law fit. Since all the distributions have the same number of free parameters, we always consider a better fit the distribution characterized by the largest normalized log-likelihood. The percentage of data samples for which a power law fits better than the exponential and Fisher log series and the corresponding exponents are presented in Supplementary Tables 1 and 2.

### OTU Analysis

We analyze metabarcoding data from marine (*33*) and freshwater (*47*) protist planktonic communities.

We retrieve the dataset of oceanic samples from European Nucleotide Archive (accession id PR-JEB16766). The dataset consists of assembled paired-end Illumina HiSeq2000 sequencing reads of PCR-amplified V9 loop of protist 18S rRNA gene obtained from 121 seawater locations distributed worldwide. In the first analytical procedure, we trim the primer sites using USEARCH (v.11.0.667) (*67*). Primer sites include 15 and 20 nucleotide sites for the 5-end and 3-end, respectively. The trimmed sequences are quality filtered with USEARCH using the option –fastq_maxee 1.0, which discards sequences with > 1 total expected errors in the sequence. The sequences are dereplicated and singleton sequences (i.e. sequences with single occurence) removed using VSEARCH (v.2.10.1) (*68*). Chimeric sequences are detected and removed using UCHIME (implemented in VSEARCH) (*69*) and a combination of reference-based (with non-redundant SILVA SSU Ref database ver.132 used as reference) and de-novo methods. Sequences are then clustered into operational taxonomic units (OTUs) using VSEARCH and the –cluster_size option. We use sequence identity thresholds of 95% and 97% which provides different levels of taxonomic resolution.

We obtain the freshwater dataset, consisting of paired-end Illumina HiSeq2500 reads of amplified genomic region encompassing V9 loop of 18S rRNA gene and ITS1 gene for 217 European freshwater lakes, from Short Read Archive (Bioproject ID PRJNA414052). First, reads from PCR replicates and sequencing replicates are merged for each lake sample. Next, primer regions are trimmed with CUTADAPT (*70*), discarding reads missing one or both of the primer sites. Forward and reverse reads with minimal overlap of 70 base pairs and with maximum of 5 nucleotide differences in the overlapping region are merged with VSEARCH (command –fastq_mergepairs). Next, we extract from the amplified SSU V9 + ITS1 region the SSU V9 region using ITSx (v.1.1.1) (*71*). This step allows the taxonomic resolution of the clustered freshwater planktonic community OTUs to closely resemble the taxonomic community resolution of the marine planktonic community, which is based on sequenced V9 loop regions of 18S rRNA genes. The extracted reads are quality-filtered, dereplicated, singletons and chimeras are removed, and the quality-filtered reads are clustered into OTUs as described above for the marine dataset. The taxonomy is assigned against SILVA v123 eukaryotic 18S subset database. OTUs assigned to Fungi, Metazoa, or Embryophyta (i.e. non-protist eukaryotes) with at least BS > 0.8 support are excluded from the final OTU tables.

## Acknowledgements

We thank M. Cencini, E. Economo, M.A. Muñoz, and L. Peliti for comments on a preliminary version of the manuscript.

## Author Contributions

P.V.M. and S.P. conceived the research; P.V.M. and S.P. wrote the code and performed the numerical simulations; A.B. and T.B. analyzed the metagenomic data; P.V.M. and S.P wrote the first draft, with subsequent input from A.B. and T.B.

